# MobileLAMP: A portable, low-cost, open-source device for isothermal nucleic acid amplification

**DOI:** 10.1101/2024.02.13.580127

**Authors:** Mohini Bhupathi, Smitha Hegde, Ganga Chinna Rao Devarapu, Jennifer C Molloy

## Abstract

Isothermal amplification-based methods for pathogen DNA or RNA detection offer high sensitivity, rapid detection, and the potential for deployment in remote fields and home testing. Consequently, they are emerging as alternatives to PCR and saw a surge in research activity and deployment for the rapid detection of SARS-CoV-2 during the Covid-19 pandemic. The most common isothermal DNA detection methods rely on minimal reagents for DNA amplification and simple hardware that can maintain isothermal conditions and read-out a fluorescent or colorimetric signal. Many researchers globally are working on improving these components based on diverse end-user needs. In this work, we have recognized the need for an open-source hardware device for isothermal amplification, composed of off-the-shelf components that are easily accessible in any part of the world, is easily manufacturable in a distributed and scalable way using 3D printing, and that can be powered using a wide diversity of batteries and power sources. We demonstrate the easy assembly of our device design and demonstrate its efficacy using colorimetric LAMP for both RNA and DNA targets.

## 2 Introduction

Isothermal amplification has emerged as an alternative to PCR for nucleic acid-based diagnostics over the last two decades. Since then, loop-mediated isothermal amplification (LAMP) (1), recombinase polymerase amplification (RPA) (2), nucleic acid sequence based amplification (NASBA) (3), strand displacement amplification (SDA), helicase dependent amplification (HDA) etc have been subject to significant innovation and increasing popularity. As of 2019, LAMP is the most published isothermal method based on Web of Science data (4). Its simple principle, high sensitivity, rapid results for diagnostic applications, dependence on a single DNA polymerase enzyme with minimal assay components has made it a popular choice for Nucleic-acid Amplification Testing (NAT) despite drawbacks due to the complexity of primer design, propensity for non-specific amplification and high risk of amplicon contamination.

Isothermal NAT such as LAMP also allow the use of simple and low-cost devices that maintain isothermal conditions, without the need for temperature cycling. Amplification has most often been monitored by fluorescence detection via DNA intercalating dyes and fluorophore-containing probes. LAMP detection methods have extended to visual readouts; without the need for any machines or lateral-flow devices (5). These simplifications offer promising applications for home-testing, field-testing or use in resource-constrained point-of-care testing. As a result of this flexibility, several LAMP assays have been commercialised for COVID-19 (6), and based on these successes several other LAMP assays are being developed for malaria (US-LAMP) (7), HIV (8) and tuberculosis (TB-LAMP) (9).

One limitation for the proliferation of LAMP outside of medical laboratories is the requirement for an incubation device that maintains a constant temperature (10). Therefore, several efforts have focused on developing a simple incubator. These efforts can be classified into two groups: instrument-free and instrument-based. Instrument-free incubators employ heating sources that don’t require electric power. Most of these instrument-free approaches consist of boiled water-baths (11–14) or phase-change materials (8,10,15–20) to provide relatively constant temperatures. However, instrument-free incubation for LAMP has several limitations, such as inability to precisely control temperature, lack of reproducibility, and potential for contamination, making it difficult to obtain consistent and reliable results.

Instrument-based incubators, on the other hand, generate constant heating by passing current through resistors such as copper and nichrome (21–23). However, most examples are either expensive or involve many complex skills, tools and materials to build. The main complexity comes from the requirement of a microcontroller to maintain a constant temperature using a proportional–integral–derivative (PID) controlling program (23,24). Another challenging part is finding a suitable heat block to hold the LAMP reaction tubes (usually 0.2 ml microtubes) and distribute the heat evenly. Additionally, these electrically incubation devices require reliable power supply, usually either AC power (25–28) or 12V DC (29–32), constraining the wide deployment of these LAMP devices.

To overcome these challenges, we have developed an incubation device for LAMP assays that is portable, affordable (<$5), robust, and easy to build. We call this incubation device “MobileLAMP” as it can be powered using widely available 5V USB power outlets, including mobile phones. In the article, we will describe the building blocks of the MobileLAMP, its fabrication and operation and evaluate its performance by demonstrating the detection of SARS-CoV-2 and *Salmonella enterica* serovar Typhi. We also discuss the benefits of MobileLAMP in comparison with existing incubator designs, such as portability, safety, and scalability. We share all the design files for MobileLAMP openly and believe that MobileLAMP can be widely adopted for other isothermal nucleic acid amplification assays.

## 3 Materials and Methods

### 3.1 Design

As shown in Figure 1, MobileLAMP is a simple and portable design consisting of a few components that can be grouped into three categories: i) electrical ii) mechanical iii) 3D-printed.

**Figure 1:**
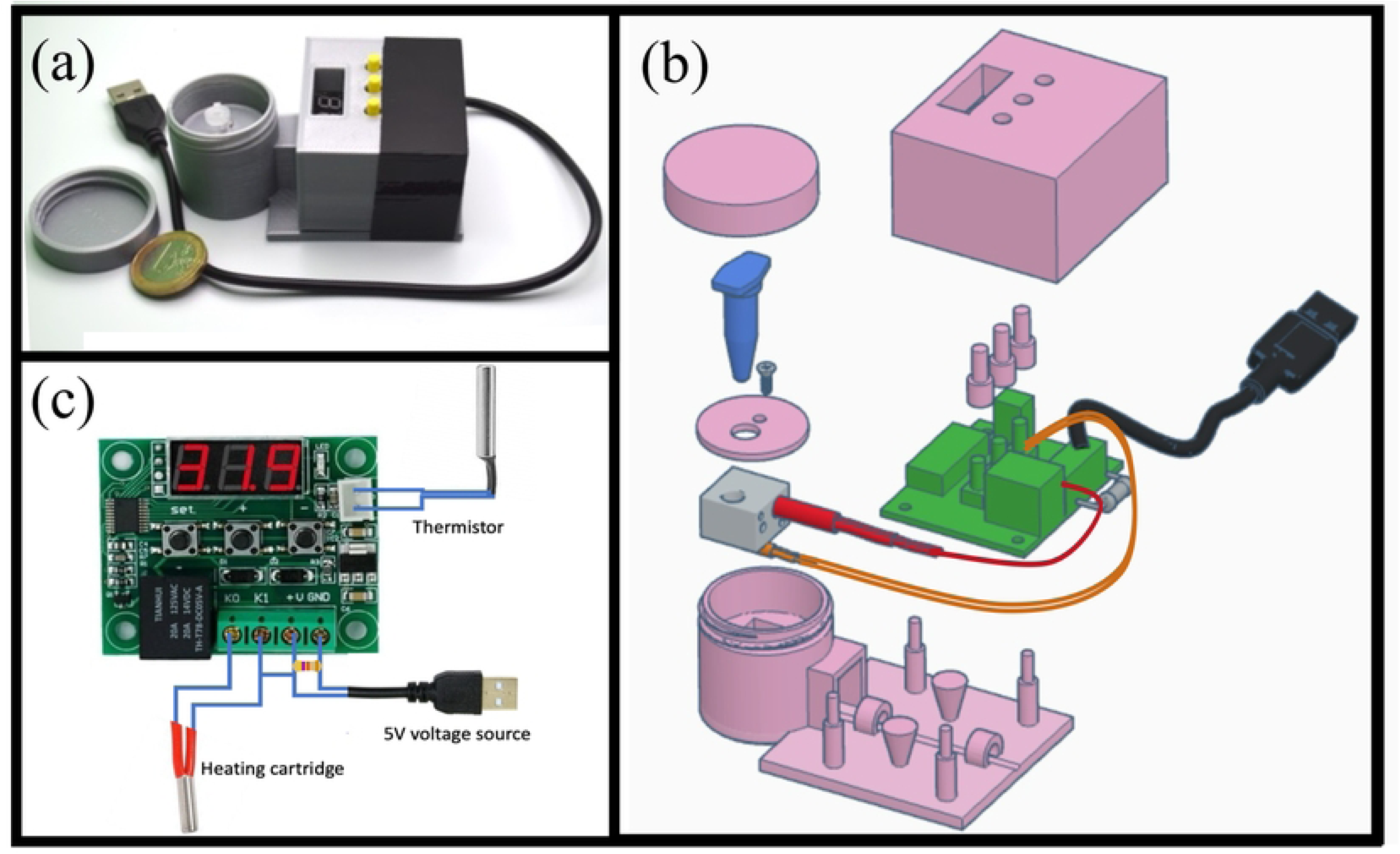
MobileLAMP design and final device image. (a) Photograph of the MobileLAMP device with a one Euro coin included for size reference. (b) Exploded view of the computer-aided design (CAD) diagram illustrating the various components of the MobileLAMP. (c) Electrical schematic of the MobileLAMP, featuring the W1209 temperature controller module.

#### 3.1.1 Electrical components of the MobileLAMP

The MobileLAMP device utilises a commercially available, low-cost ($1.5) thermostat module known as W1209 (Figure 2a). W1209 module is a no-code programmable thermostat that comprises three tactile switches, and a seven-segment LED display, enabling configuration of various parameters to set a desired temperature in the range of –50 °Cto 110 °C (see supplementary section A for more information). As a result, the user does not need any programming knowledge to operate the MobileLAMP device. The W1209 also comes along with a temperature sensor (Figure 2b) which provides real-time temperature monitoring and feeds information to a preconfigured proportional–integral–derivative (PID) algorithm to accurately adjust the temperature.

**Figure 2:**
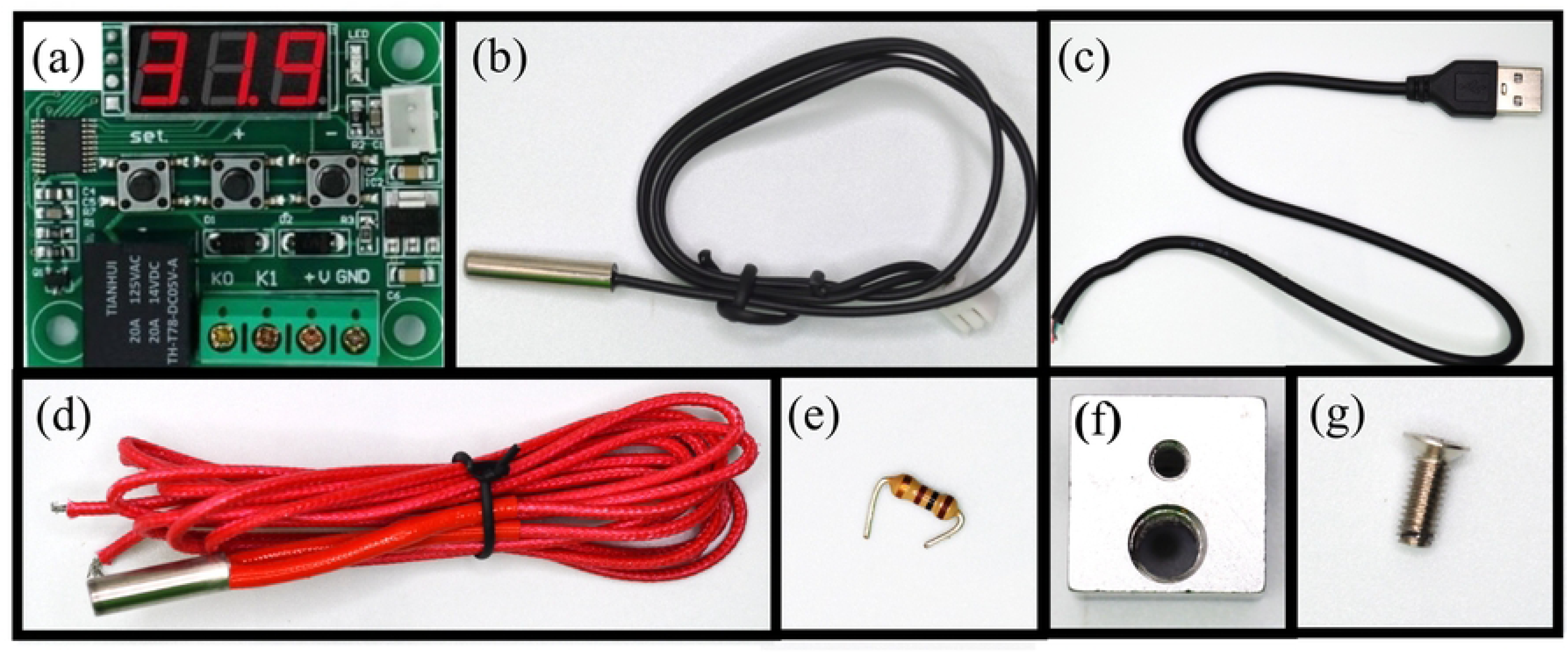
Electrical and mechanical components required for building the MobileLAMP device: (a) W1209 temperature controller module. (b) thermistor (c) USB cable (d) heater cartridge (e) 100 Ohm resistor (f) heat block (g) 3mm screw.

The W1209 module operates at 5V, making it possible to power the MobileLAMP using widely available 5V USB power sources, such as mobile phones, chargers, and power banks. To facilitate this, a USB-male cable (Figure 2c) is employed as the power cable for the MobileLAMP. To provide a stable temperature for LAMP reactions, a heating cartridge (Figure 2d) commonly used in Fused Deposition Modelling (FDM) 3D printer extruders is utilised as the heat source within the MobileLAMP. A 100 ohm resistor (Figure 2e) is used in MobileLAMP to draw the minimum required current to ensure that it works with the power banks as some rechargeable power banks require a minimum current draw to maintain a steady power supply.

#### 3.1.2 Mechanical components of the MobileLAMP

The MobileLAMP device utilises an aluminium heating block (Figure 2 (f)), commonly found in FDM 3D-printer extruders, as its heating block. This block features two 6 mm diameter holes, one for holding a heating cartridge and one for a nozzle in 3D-printer extruders. These holes are repurposed in the MobileLAMP to hold the heating cartridge and a microtube containing the LAMP reaction mixture. Due to its widespread usage, the aluminium heating block can be obtained at a low-cost of less than $1. Additionally, a 3mm screw (Figure 2 (g)) is utilised in the construction of the MobileLAMP to secure the heating cartridge in the heating block as detailed in Section 3.2.

#### 3.1.3 3D-printed parts of the MobileLAMP

The MobileLAMP consists of seven 3D-printed parts, designed using OpenScad (RRID:SCR_018870) (33), a free and open source parametric CAD software. Figure 3a to 3e show the screenshots of the CAD images of these 3D-printed parts. The bottom part of the enclosure (Figure 3a) is designed to accommodate the heat block, the temperature sensor and the W1209 thermostat electronic controller. One notable aspect of the enclosure is the jar and lid design (Figure 3c), along with a circular cover (Figure 3d) helps to minimise heat loss from the heating block to the environment. The top part of the enclosure (Figure 3b) has provision for three 3D-printed buttons (Figure 3e) which allow users to interact with the tactile switches of the W1209 and control heat block’s temperature. Additionally, the top part of the enclosure includes a window for the 7-segment LED display of the W1209 module. The CAD design files of the MobileLAMP were exported as STL files from Openscad and then converted to G-code using CURA (RRID:SCR_018898) (34), an open-source slicer program that slices 3D CAD models into 2D layers. Care was taken to print the parts without any support structures to reduce plastic waste and reduce assembly time. The G-code files were then uploaded to an Ultimaker 3 3D-printer (35) and printed using PLA material with the following settings: 2mm layer height, 20% infill and no support structures. The print orientation of the objects of the MobileLAMP are shown in Figure 3 (f). Figure 4 shows these 3D-printed parts of the MobileLAMP.

**Figure 3:**
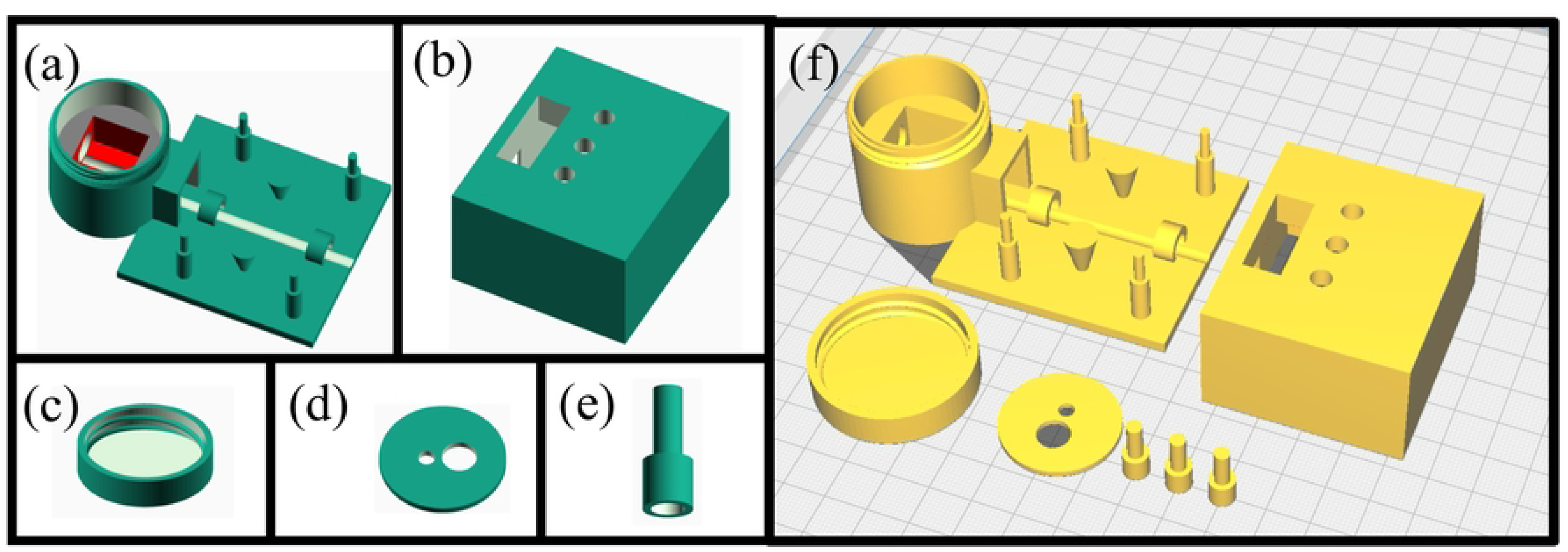
MobileLAMP Computer-aided design (CAD) diagrams. (a) to (e) CAD diagrams of the MobileLAMP’s 3D-printed components designed using OpenScad software. (f) Representation of the print orientation of the MobileLAMP’s 3D-printed parts in the CURA slicing software.

**Figure 4:**
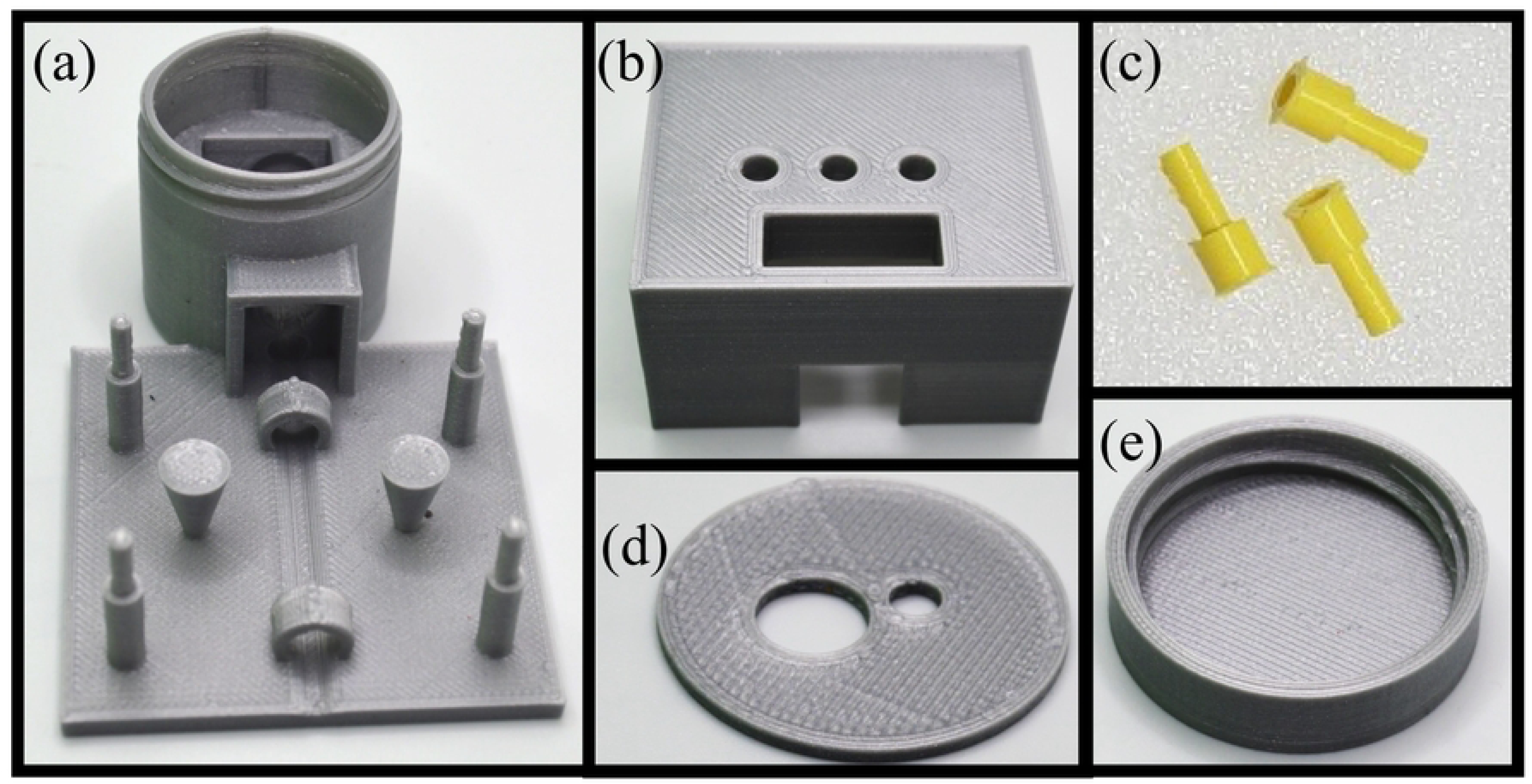
Three-dimensional (3D) printed components of the MobileLAMP using Polylactic Acid (PLA) material. (a) Bottom portion of the electronic enclosure. (b) Top portion of the electronic enclosure. (c) Tactile switch buttons for the W1209 module. (d) Circular cover for the heat block. (e) Heat chamber lid for the MobileLAMP.

#### 3.1.4 Bill Of Materials

**Table 1.** Bill of Materials to construct the MobileLAMP. Note that the cost of these 3D-printed parts is estimated assuming a $22.99 per kg spool of PLA material (**36**).

| Component | Parameter<br>s | Number | Cost (\$) | Total Cost (\$) | Source Of<br>Materials | Material<br>Type |
| --- | --- | --- | --- | --- | --- | --- |
| Electrical components |  |  |  |  |  |  |
| W1209<br>module | 5V version | 1 | 1.5 | 1.5 | AliExpress | NA |
| USB cable | Male | 1 | 0.5 | 0.5 | AliExpress | NA |
| Heating<br>cartridge | 14 Ohms | 1 | 1 | 1 | AliExpress | Ceramic |
| Resistor | 100 ohms | 1 | 0.01 | 0.01 | AliExpress | NA |
| Mechanical components |  |  |  |  |  |  |
| Heat block | E3D V5 | 1 | 1 | 1 | AliExpress | Aluminium |
| Screw | 3mm | 1 | 0.01 | 0.01 | AliExpress | NA |
| 3D-printed components |  |  |  |  |  |  |
| Top part | NA | 1 | 0.275 | 0.275 | In House | PLA |
| Bottom part | NA | 1 | 0.528 | 0.528 | In House | PLA |
| Circular<br>cover | NA | 1 | 0.0229 | 0.0229 | In House | PLA |
| Lid | NA | 1 | 0.114 | 0.114 | In House | PLA |
| Pins | NA | 3 | 0.007 | 0.02299 | In House | PLA |
| Total Cost<br>(EUR) |  |  |  | 4.98 |  |  |

### 3.2 Build instructions of the MobileLAMP

A. Begin the construction of the MobileLAMP with placing the bottom enclosure part (Figure 5a).
B. Insert the thermistor into the designated slot within the enclosure (Figure 5b).
C. Place the heat block inside the cylindrical chamber of the bottom enclosure (Figure 5c).
D. Fit the 3D-printed circular cover atop the heat block (Figure 6a).
E. Insert the heating cartridge into the heat block, and secure it by tightening the 3mm screw from the top of the circular cover (Figure 6b).
F. Mount the W1209 module to the enclosure using the four designated mounting pillars (Figure 6c).
G. Carefully follow the circuit diagram (Figure 1b) to make the necessary electrical connections (Figure 7a).
H. Insert the 3D-printed button pins into the tactile switches of the W1209 (Figure 7b).
I. Finally, fit the top part of the enclosure onto the bottom part, ensuring that the 3D-printed buttons are not obstructed (Figure 7c).
J. Secure the top and bottom parts of the enclosure together with tape to complete the construction of the MobileLAMP (Figure 8a).

### 3.3 Operational instructions of the MobileLAMP

A. To operate the MobileLAMP, connect it to a 5V USB power supply using its USB cable.
B. Set the desired temperature by adjusting the parameters P0 to P6 on the W1209 module’s settings using the 3D-printed buttons.
C. Refer to Figure S1 and S2 for detailed instructions on how to navigate the settings.
D. The W1209 module will execute the program with the selected parameters and display the real-time temperature on the seven segment LED display.
E. Wait until the MobileLAMP reaches the set temperature, as indicated on the display.
F. Once the desired temperature has been reached, place the LAMP reaction tube inside the heating block (Figure 8 (b)).
G. Secure the lid by screwing it into the reaction chamber jar (Figure 8 (c)).
H. Allow the LAMP reaction to proceed for the designated amount of time before removing the lid.
I. Carefully remove the LAMP reaction tube from the heating block and observe the tube for any colour changes, which will indicate the outcome of the LAMP reaction.

### 3.4 Colorimetric LAMP Protocol

We performed colorimetric LAMP on synthetic targets of SARS-CoV-2 and *S.* Typhi (the bacterial pathogen that causes typhoid fever) using 2X Warmstart colorimetric LAMP (M1800, NEB, US). The RNA template of the SARS-CoV-2 N gene was synthesised using *in-vitro* transcription (Hiscribe, NEB, US) and a template plasmid (Molecular Diagnostics Collection, Free Genes, Stanford University) using forward and reverse primers N-RNAF and N-RNAF (Table 2). Final working concentrations of SARS-CoV-2 primers were 1.6 µM FIP/BIP, F3/B3 0.2 µM and L3/B3 0.4 µM. *S.typhi* synthetic template targeting *STY1607* gene (37) were synthesised (Twist Bioscience, US) and the LAMP primers (Table 2) of final concentration 3.2 µM FIP, 1.6 µM BIP, F3/B3 0.2 µM and LB/LF 0.8 µM were used. 10 µl of the reaction mixture at various target concentrations along with a no DNA target blank (NTC) were incubated in MobileLAMP for 60 minutes with quick spinning every 15 minutes in a mini centrifuge to reduce any collection of condensate at the lid. These reactions were also incubated using a thermocycler (miniPCR bio™, US), for comparison.

**Figure 5:**
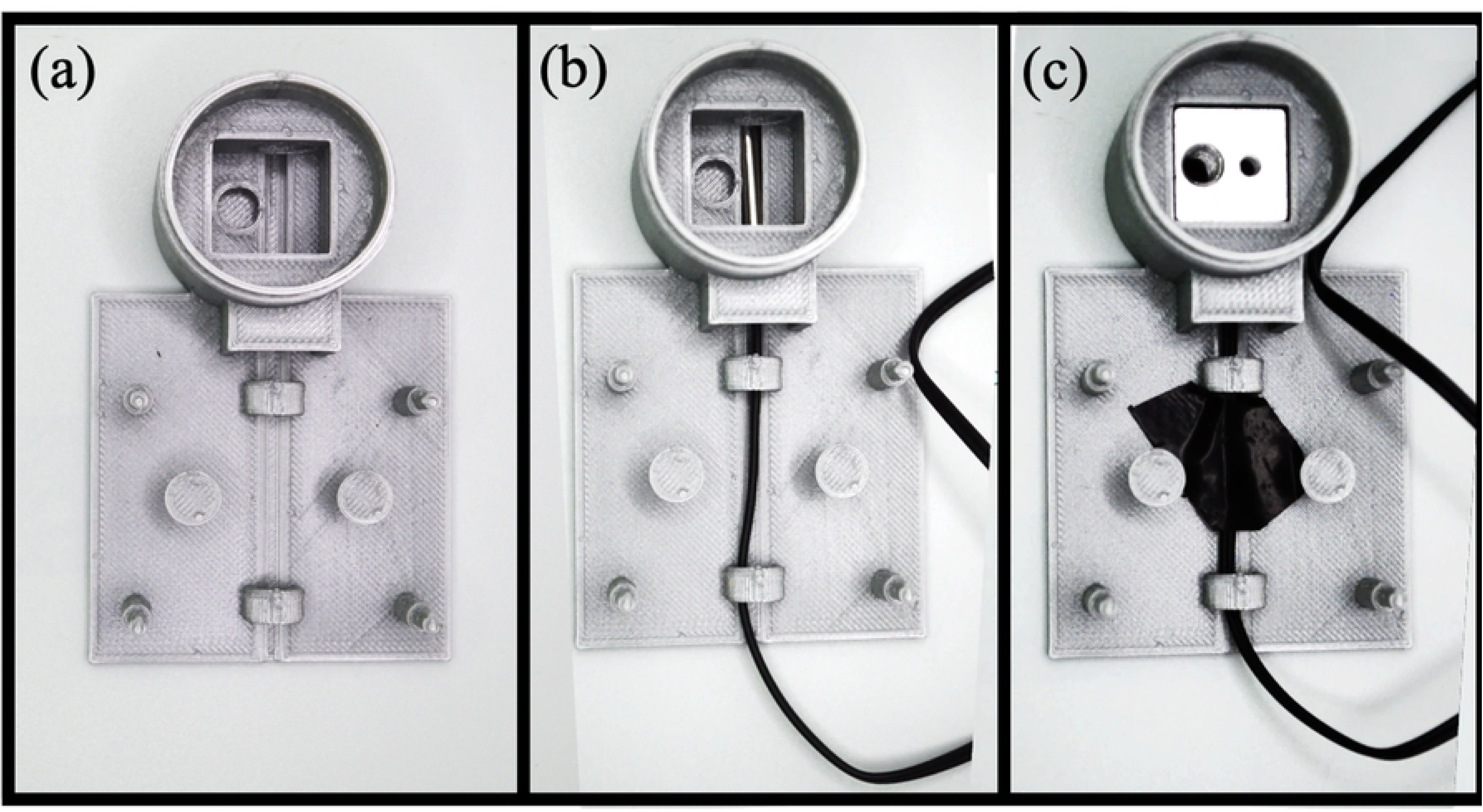
Assembling MobileLAMP enclosure and thermistor. (a) 3D printed part of MobileLAMP’s bottom enclosure. (b) Thermistor is inserted into the slot. (c) Heat block is placed inside the cylindrical chamber of the bottom enclosure. Thermistor wire is held in place using a black tape.

**Figure 6:**
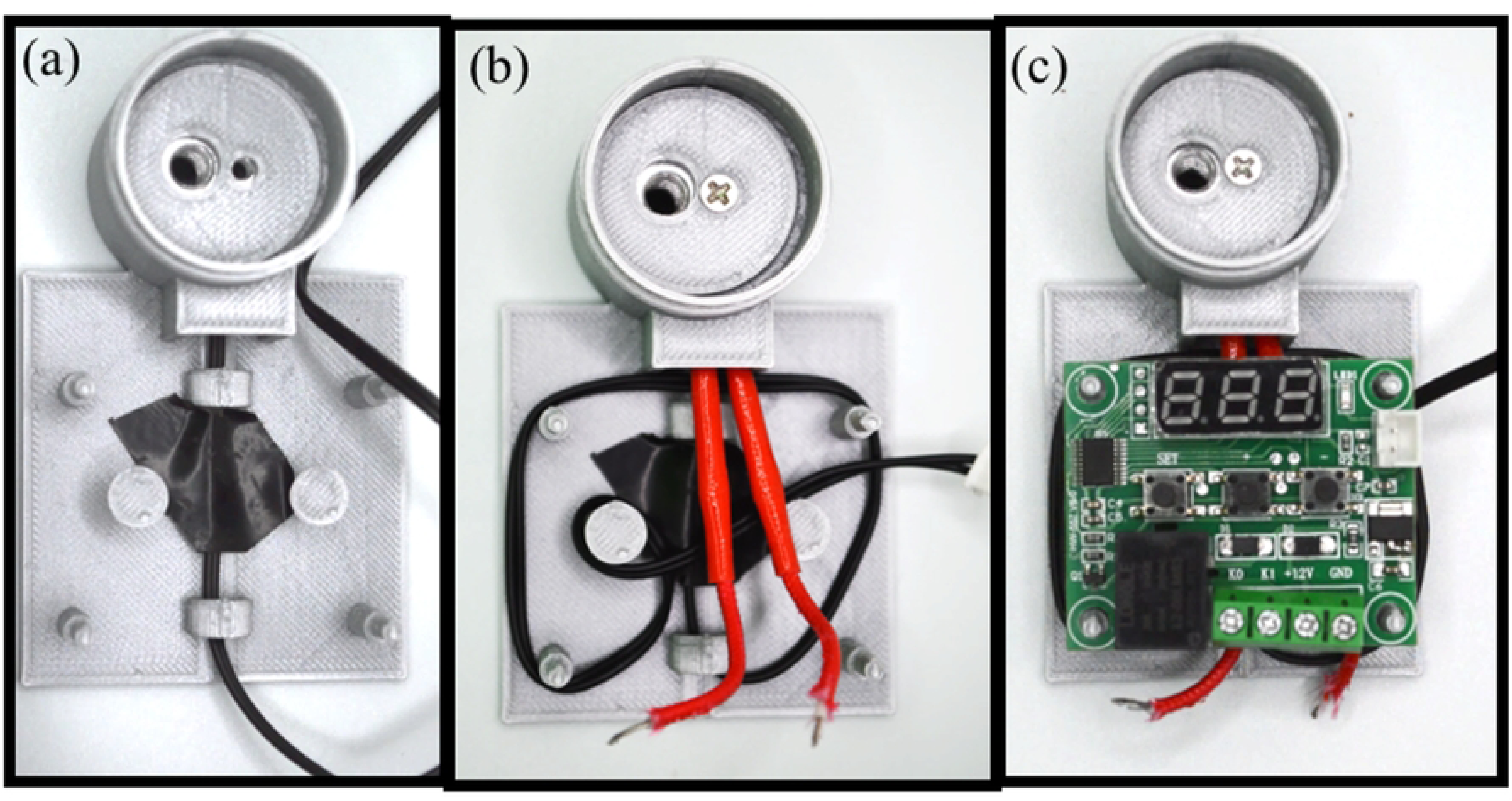
Assembling MobileLAMP heating components. (a) A 3D-printed circular cover is fitted atop the heat block (b) The heating cartridge is inserted into the heat block and secured in place with a 3mm screw (c) The W1209 module is mounted to the enclosure using the mounting pillars.

**Figure 7:**
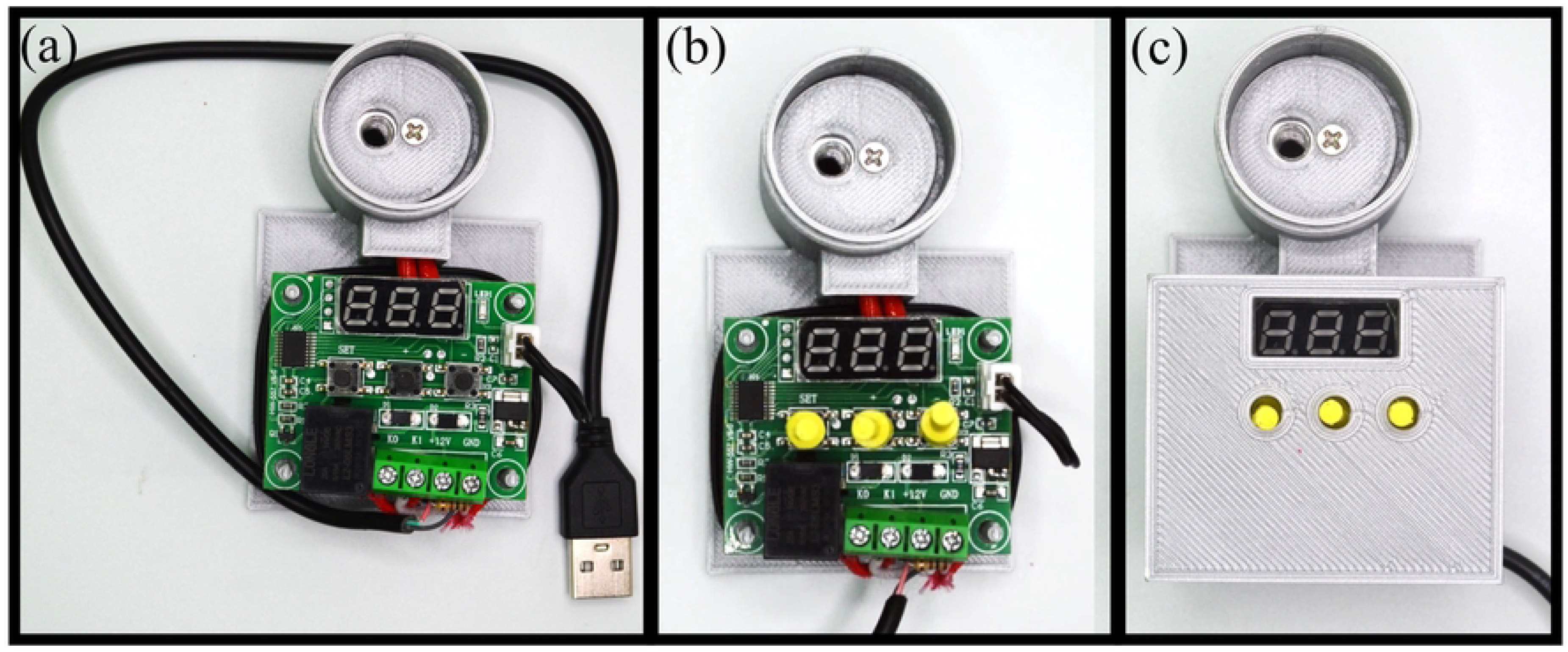
Assembling MobileLAMP electrical components. (a) Complete the electric connections according to the circuit diagram shown in Figure 3. (b) Insert the 3D-printed button pins into the tactile switches of the W1209. (c) Secure the top cover to the enclosure.

**Figure 8:**
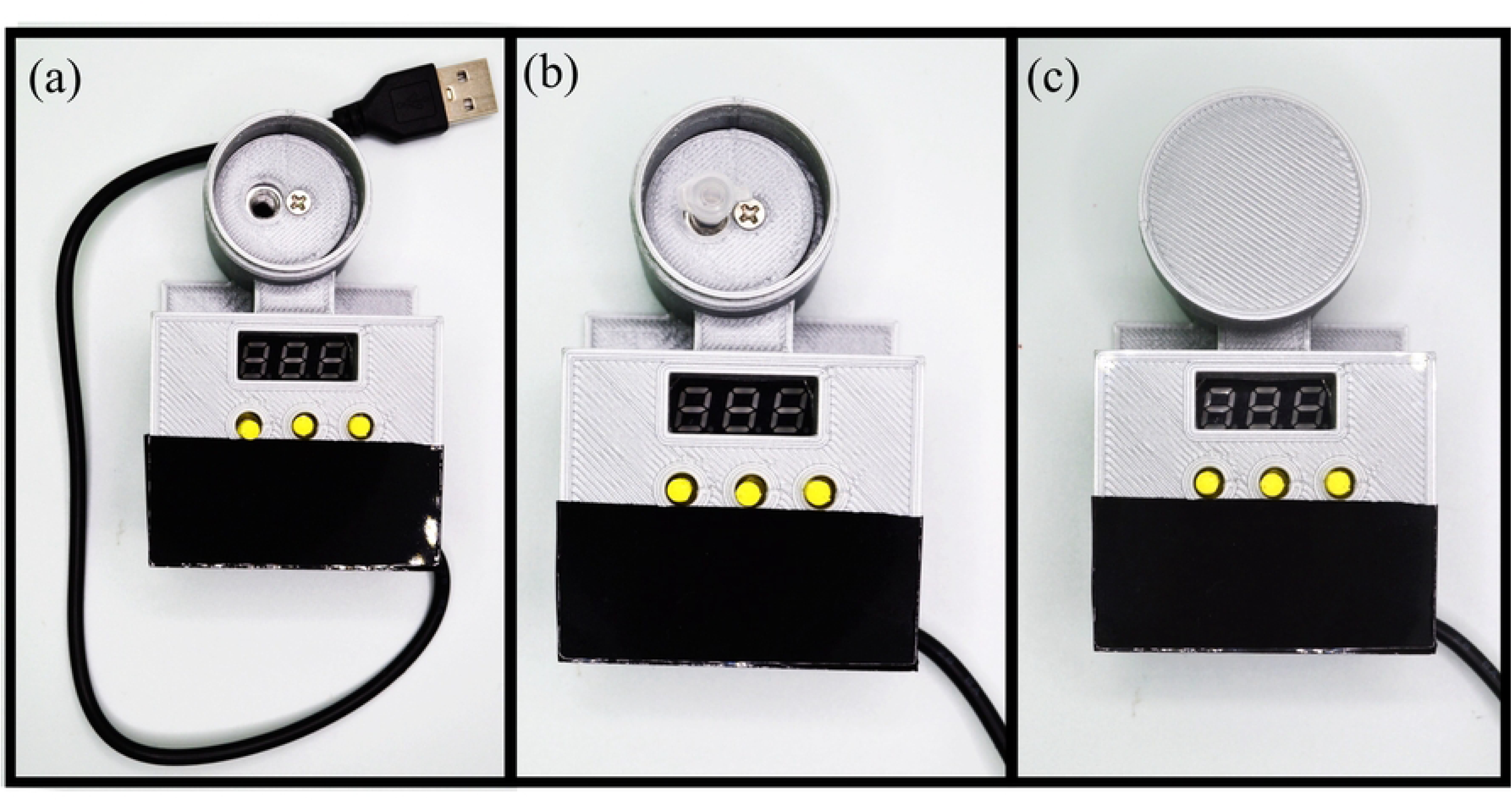
Final assembly steps of MobileLAMP. (a) The top and bottom parts of the lamp device are secured with tape. (b) The lamp reaction tube is inserted into the heat block. (c) The upper lid part is fixed to the reaction chamber of the device.

**Table 2:**
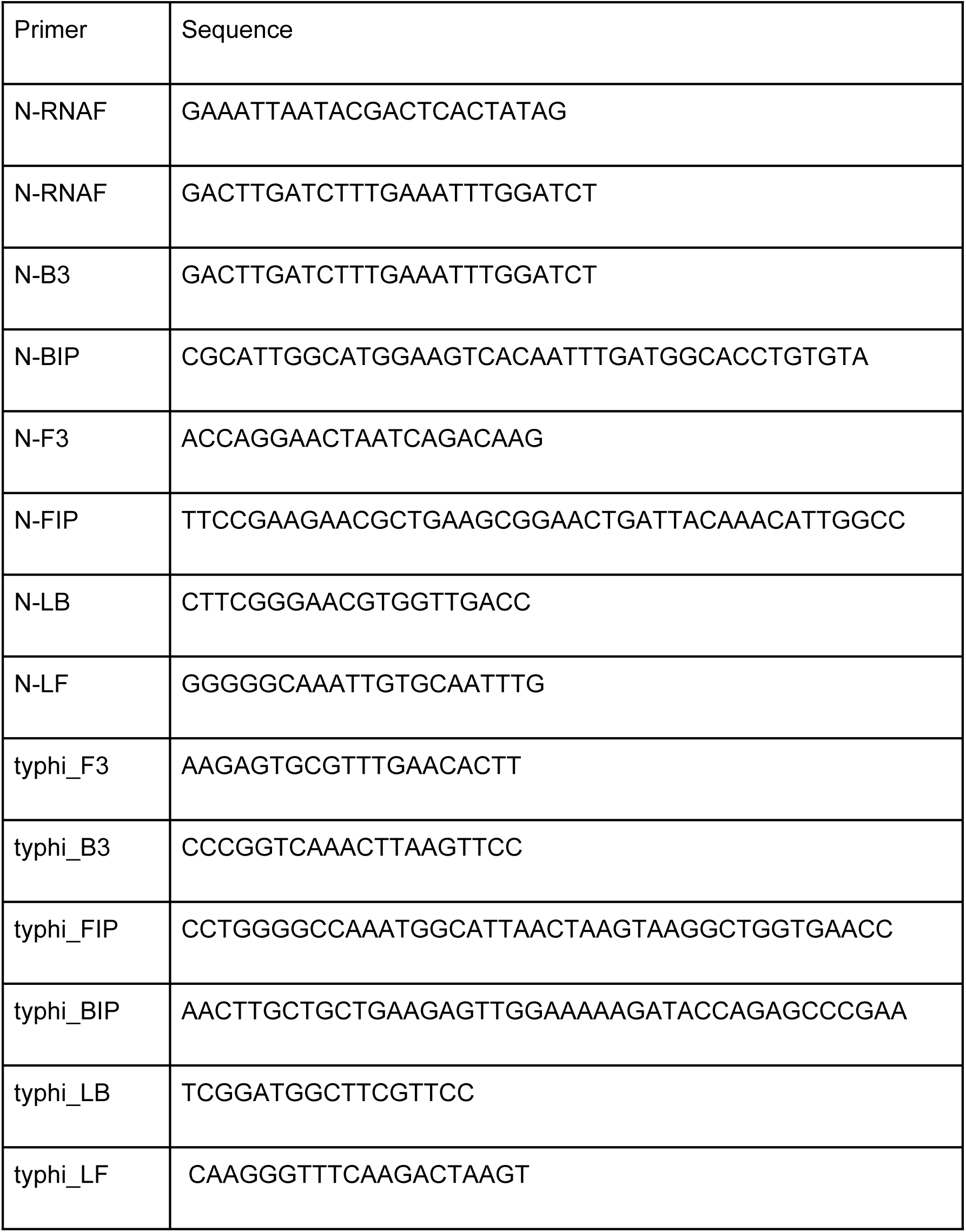
List of primers used in colorimetric LAMP for the detection of SARS-CoV-2 and *S.typhi*.

## 4 Results

### 4.1 Temperature characterization

In this work, a temperature of 65°C is used for all LAMP reaction experiments. To achieve this temperature, the W1209 module is set to the following parameters: Target temperature set to 62.5°C, P0 set to H, P1 set to 0.5°C, P2 set to 62.5°C, P3 set to 65°C, P4 set to 62°C, P5 set to 7°C and P6 set to OFF. To validate the MobileLAMP performance, we additionally evaluated the stability of the set-point temperatures using a 10kΩ commercially available NTC thermistor (Vishay, USA) using an Arduino Uno microcontroller. The thermistor was immersed in a microtube with water and the temperature profile at 65°C was monitored for 60 minutes as shown in Figure 9 (a). The mean temperature of three repeats for an hour was measured to be 64.7 ℃ and a variance of 0.2℃.

**Figure 9:**
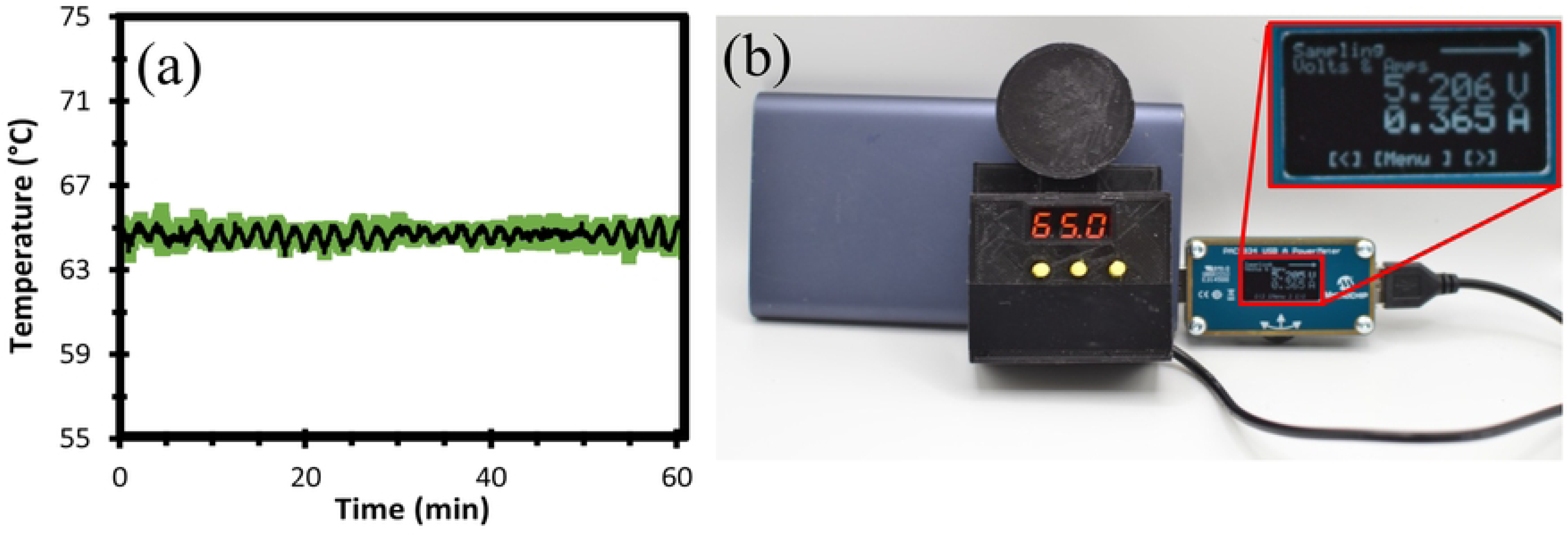
MobileLAMP temperature and power characterisation. (a) MobileLAMP temperature profile at 65 °C set-point. Black line is the mean temperature value (three repeats), green shaded region denotes one standard deviation from the mean (b) The power consumption of the MobileLAMP device is measured with USB-A power meter (PAC1934 from Microchip Technology Incorporated), by connecting the power meter between the USB power source and the MobileLAMP. Current consumption of the MobileLAMP (365 mA) is shown in the inset.

### 4.2 Electrical characterization

We measured the power consumption of MobileLAMP using a USB-A power meter (PAC1934 USB-A from Microchip Technology Incorporated). The test setup involved connecting the power meter between the USB source and the MobileLAMP device, as shown in Figure 9 (b). It is evident from Figure 9 (b) that the MobileLAMP device draws only 365 mA of current which falls comfortably below the standard 500 mA provided by computers, mobile phones, and comparable devices for connecting USB peripherals.

### 4.3 Colorimetric LAMP against various targets

We first performed the colorimetric RT-LAMP to detect SARS-CoV-2 RNA with the MobileLAMP and compared it with a thermocycler as shown in (Figure 10). The measurements were done in duplicate at three different target concentrations (602, 6022 and 60221 copies/µl) compared against the NTC. The reactions turned yellow from pink when the target was detected and amplified in both the devices. We next performed colorimetric LAMP assay with the MobileLAMP to successfully detect with *S.typhi* gene *STY1607* DNA at a concentration of 60221 copies/µl as shown in Figure 11.

**Figure 10:**
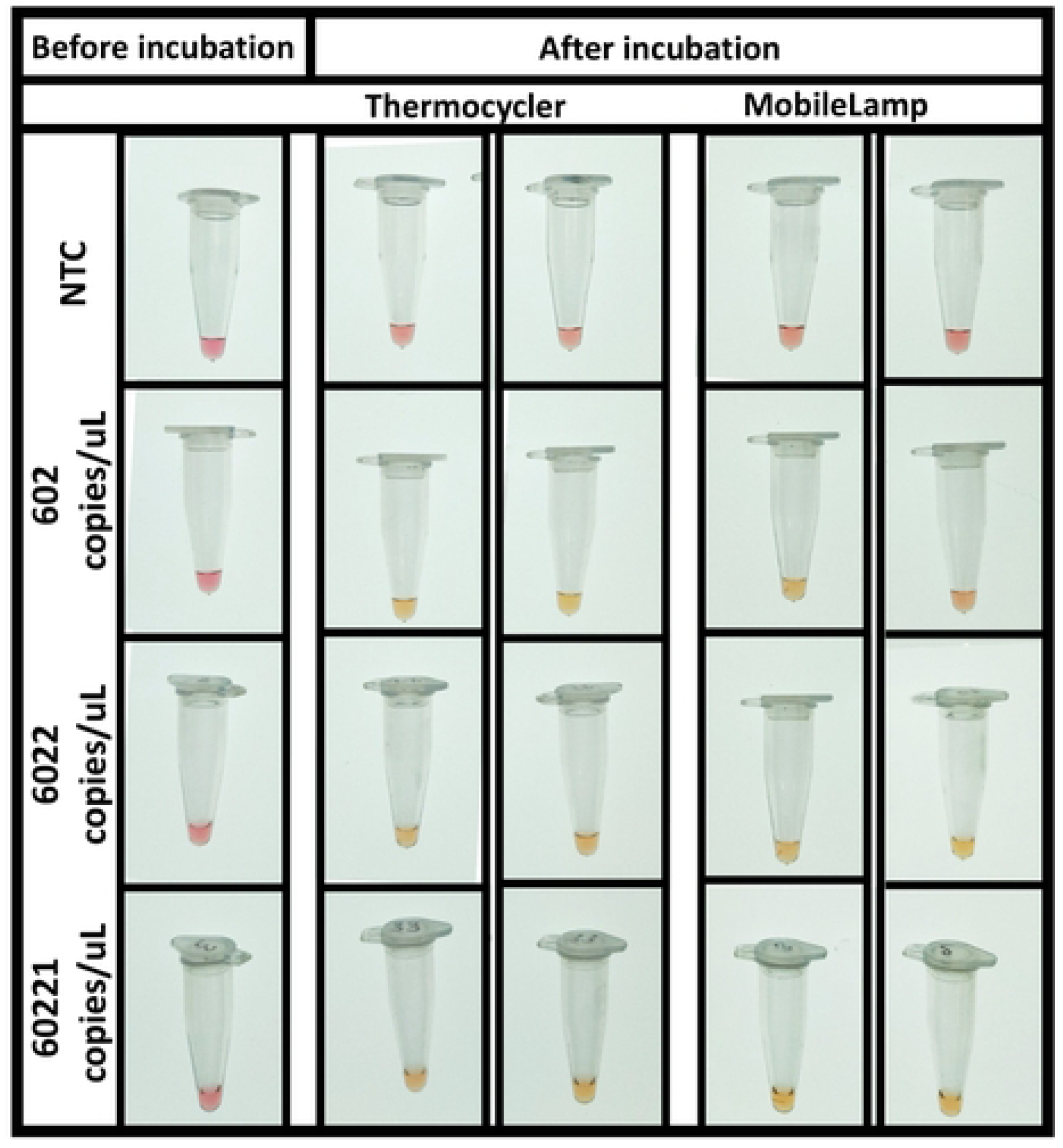
Detection of SARS-CoV-2 target using colorimetric RT-LAMP using MobileLAMP compared with thermocycler. The detection was carried out at 602 copies/ul (second row),6022 copies/µl (third row) and 60221 copies/ul (fourth row) compared to NTC. MobileLAMP successfully amplified the targets as indicated by tubes turning pink to yellow.

**Figure 11:**
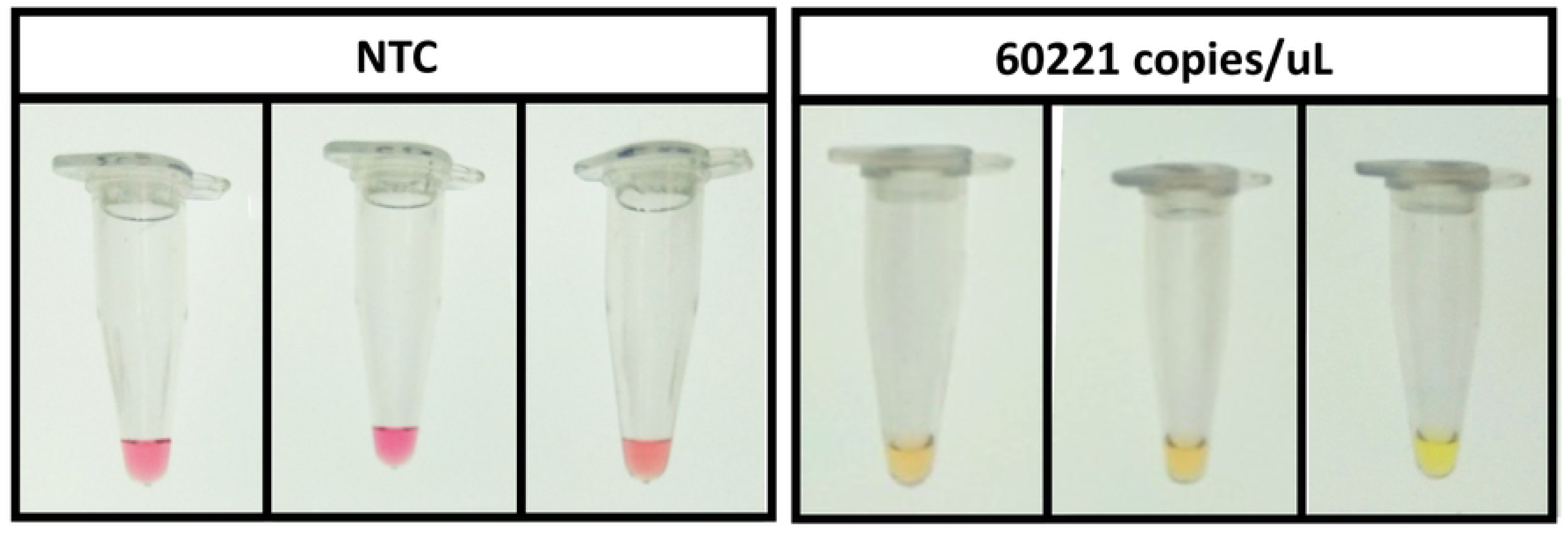
MobileLAMP for *S.typhi* detection: NTC (left) showed no colour change, while the target *STY1607* gene at 60221 copies/µl (right) changed colour to yellow indicating successful amplification.

## 5 Discussion

### 5.1 Advantages of the MobileLAMP

The MobileLAMP device provides several advantages compared to the existing similar devices in the field of isothermal nucleic acid amplification.

#### 5.1.1 Affordability

The bulk of the cost associated with building low-cost LAMP devices often comes from the electronic controller and heat block components. The MobileLAMP reduces these costs by utilising off-the-shelf components that are both low-cost and widely available. The W1209 module used in the MobileLAMP is an affordable ($1.5) temperature controller that can be easily obtained even in single quantities. Note that the W1209 module was used in a LAMP incubation device recently (38), however does not include detailed working of the module which we have covered in this work. The heat block used in MobileLAMP is a widely used aluminium block in FDM 3D-printers, which can be purchased for as little as $1 even in low quantities. This results in a total cost of the MobileLAMP being less than $5 including 3D-printed parts, making it the lowest-cost instrument-based LAMP incubator available, and even lower cost than some instrument-free LAMP incubators.

#### 5.1.2 Simplicity

The MobileLAMP is designed with a minimum number of components, making it quick and easy to build, even for non-experts. In comparison, many instrument-based LAMP devices in the literature (Ganguli et al., 2020; Papadakis et al., 2022; Buultjens et al., 2021; Velders et al., 2018, Das et al., 2023; Myers et al., 2022; McCloskey et al., 2022; Torezin Mendonça et al., 2022; Vural Kaymaz et al., 2022) require a wide range of skills, including programming, soldering, microfluidics, CNC routing and 3D-printing, making them difficult to build. This simplicity in design makes it applicable in (i) resource and budget constrained POCs where multiple units of MobileLAMP can be set-up depending on the patient load and (ii) integrated into home-testing kits with colorimetric LAMP.

#### 5.1.3 Scalability

The design of the MobileLAMP is low-cost and open-source, aiding scalability of supply. While it uses technologies such as FDM 3D-printing that are not well-suited to centralised mass manufacturing, it is possible to produce in large quantities, if required, with the help of the 3D printing community through distributed manufacturing. This approach was prevalent during the COVID pandemic for PPE manufacture, and has been explored as a way to increase access to medical devices. The small size of MobileLAMP also makes it possible to print copies on low-cost 3D printers, or to print multiple units simultaneously on large 3D-printers. Its open-source design enables users to modify and customise the device to meet their specific needs, which can foster innovation and collaboration within the scientific community.

#### 5.1.4 Deployability

MobileLAMP operates at 5V, requiring only 365mA of current, which can be supplied by ubiquitous 5V USB powered devices, such as mobile phones, and associated power supplies, such as battery banks, which are widely available. For example, the MobileLAMP can run on a 10000mAh power pack for 27 hours, or the equivalent of 27 assays. This low-power requirement along with its portability makes the MobileLAMP appropriate for field use, which can increase the efficiency and speed of sample collection and analysis. It also provides the ability to perform analysis in remote locations, where laboratory and diagnostic infrastructure may be limited.

### 5.2 Limitations of MobileLAMP

#### 5.2.1 Limited number of tubes

One of the limitations of the MobileLAMP device is its limited capacity. It can only hold one reaction tube at a time, which was a necessary trade-off to use the off-the-shelf heat block. While one sample is adequate for the intended application of the device, it does not allow for a negative or positive control sample to be run simultaneously. However, it is worth noting that even commercial LAMP devices (6) designed for Point-of-Care (PoC) applications typically have provision for only one tube, so this limitation is not unique to the MobileLAMP device.

#### 5.2.2 Lack of heated lid

Another limitation of the MobileLAMP device is its lack of a heated lid on top of the reaction chamber, which is commonly used in commercial LAMP devices to prevent evaporation of the reaction mixture and to maintain uniform mixing of the reagents with the sample. While the simplified design of the MobileLAMP device without the heated lid offers advantages in terms of ease of use and cost, this lack of lid can result in evaporation and non-uniform mixing of the reagents. This can be mitigated by using mineral oil on top of the reaction mixture, by using a larger volume of reaction mixture or, less conveniently, by periodically centrifuging the tube.

## Conclusions

MobileLAMP provides the highest level of functionality that we have observed for a LAMP device with an equivalent part cost of <$5. It has the additional advantage of simple assembly, achieved through repurposing off-the-shelf components and leveraging 3D printing to produce a device that has favourable technical characteristics for isothermal amplification at point of care. MobileLAMP can hold an average temperature of 64.7℃ with a variance of 0.2℃ over the course of an hour, and can run on a 10000mAh power pack for 27 hours, or the equivalent of 27 assays. The trade-offs include being limited to a single tube capacity and lack of heated lid. However, neither of these preclude its use to successfully run LAMP reactions for field research and educational use, and it is amenable to distributed mass manufacture in a crisis situation, contingent on the continuation of supply chains for consumer electronics. The designs have been made available under an Open Hardware License to encourage replication and improvement by other researchers.

## Acknowledgments

JM was supported by a Shuttleworth Foundation Fellowship and the Isaac Newton Trust. SH was funded by University of Cambridge EPSRC GCRF Impact Acceleration Account. The funders had no role in study design, data collection and analysis, decision to publish, or preparation of the manuscript.

## Supplementary Material

**Figure S1:**
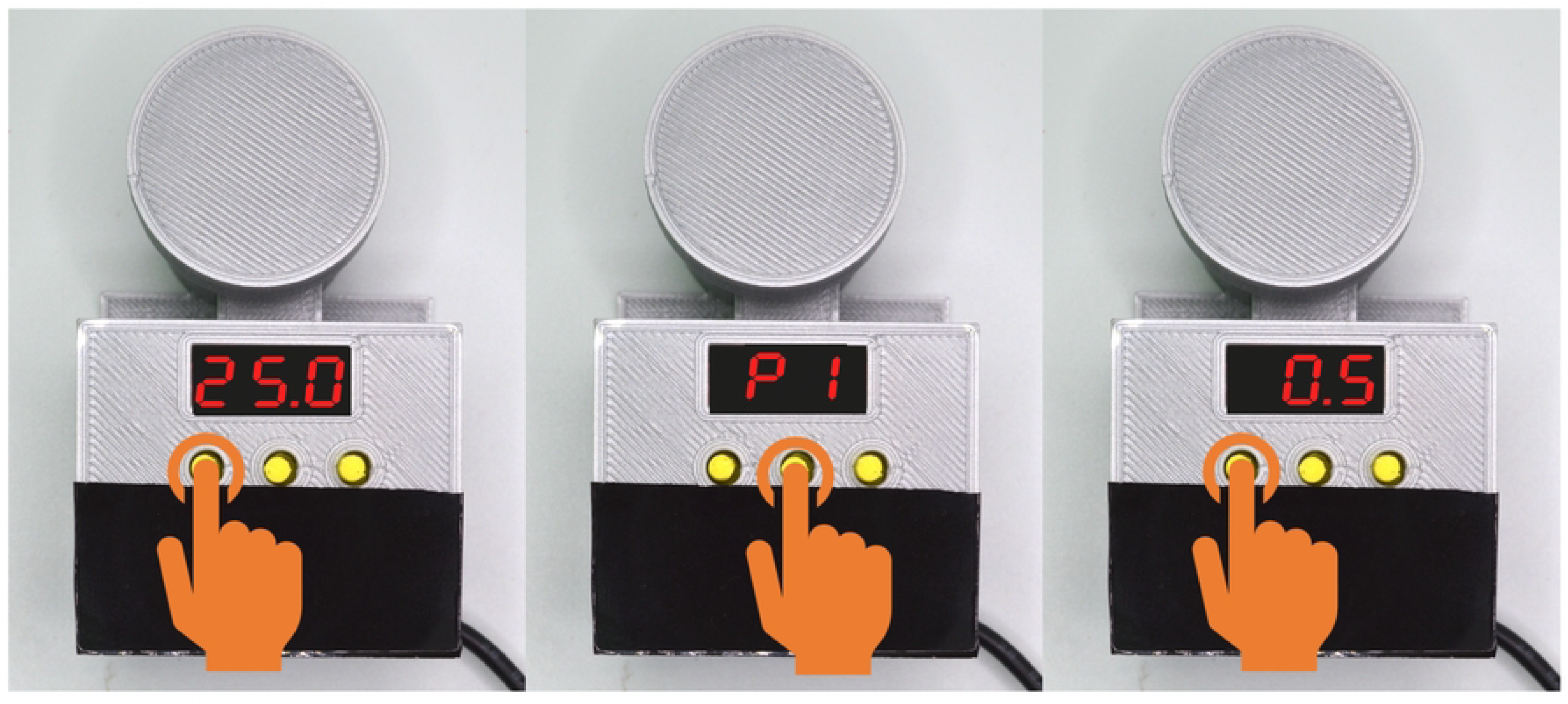
Operating instructions for setting temperature on MobileLAMP device. To set a target temperature on the MobileLAMP device, follow these steps: (a) Press the left button once to enter into the temperature change mode. (b) Use the middle button to increase the temperature. (c) Use the right button to decrease the temperature.

**Figure S2:**
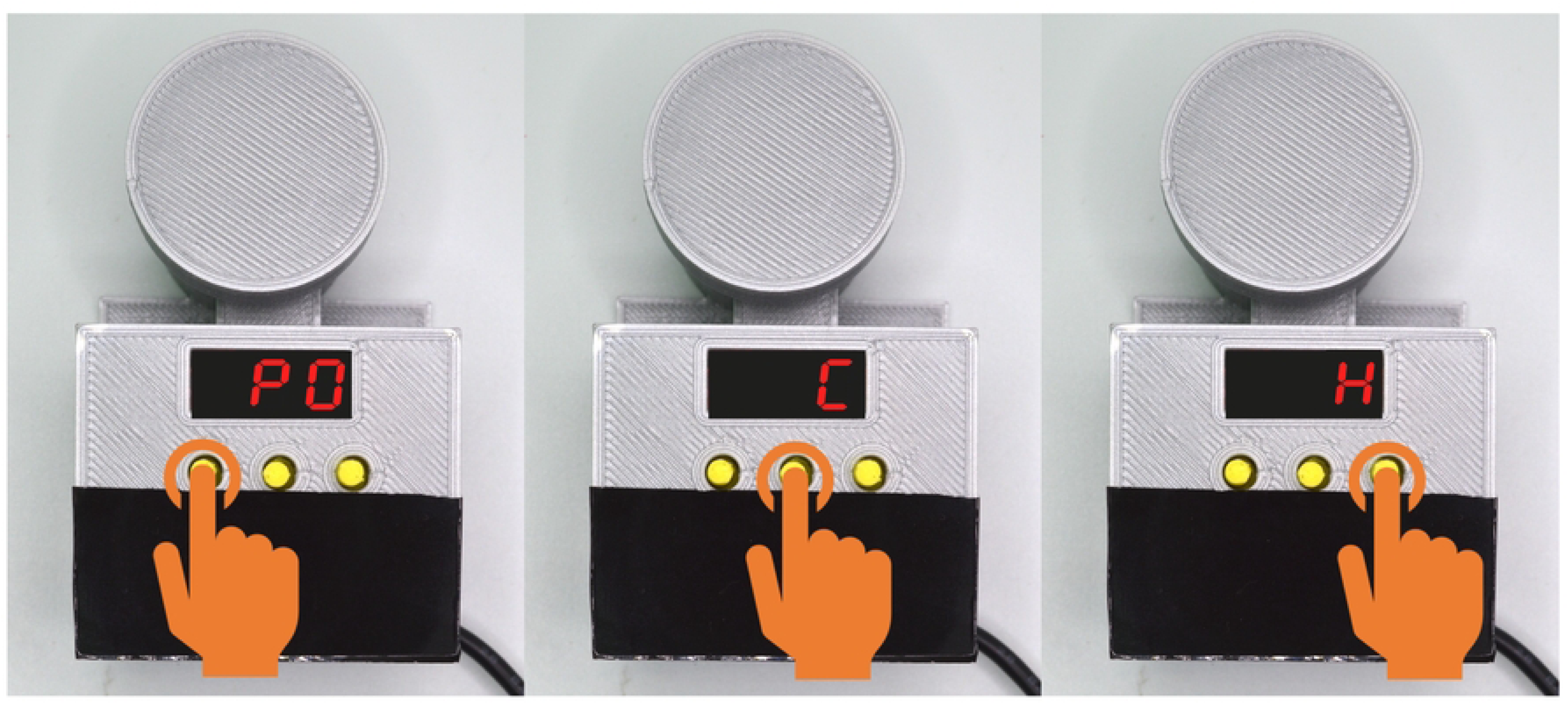
Operating instructions for additional settings on MobileLAMP device. (a) To access the additional settings of the MobileLAMP device, press and hold the left button. (b) Use the middle and right buttons to navigate through the settings menu. (c) Once the desired setting is identified, press the left button to enter it, and then use the middle and right buttons to adjust the value of the selected setting. Press the left button to exit the menu.

**Table S1.** Various settings of the W1209 module with the settings ranges and default values.

